# Design and Evaluation of an Affordable Hypoxic Chamber with Comprehensive Environmental Control for Research Applications

**DOI:** 10.1101/2023.06.21.546032

**Authors:** Syed Aasish Roshan, Swaminathan K Jayachandran, Mahesh Kandasamy, Muthuswamy Anusuyadevi

**Affiliations:** Molecular Neuro-Gerontology Laboratory, Department of Biochemistry, School of Life Sciences, Bharathidasan University, Tiruchirappalli-620024, Tamil Nadu, India; Drug Discovery and Molecular Cardiology Laboratory, Department of Bioinformatics, School of Life Sciences, Bharathidasan University, Tiruchirappalli-620024, Tamil Nadu, India; Laboratory of Stem Cells and Neuroregeneration, Department of Animal Science, School of Life Sciences, Bharathidasan University, Tiruchirappalli -620024, Tamil Nadu, India; University Grants Commission-Faculty Recharge Program (UGC-FRP), New Delhi 110002, India

**Keywords:** Invivo, CO2 adsorber, Arduino controller, Hypoxia chamber, humidity, temperature

## Abstract

**Purpose:** In-vivo Hypoxia chamber is sought after for research on sleep apnea, among others. Recently hypoxia chambers are utilized to create intermittent hypoxia, thereby utilizing it as a treatment strategy. The commonly used Invivo hypoxic chambers only monitor oxygen levels and fail to read up on vital factors like CO2 buildup, and humidity among others. In addition, these devices are expensive and make them unaffordable for many labs.

**Methods:** Here we report a lab-designed and assembled Arduino microcontroller-based hypoxic chamber with automatic maintenance of O2 level, CO2 level, and Humidity. The entire 650$ setup is inclusive of an Arduino module with sensors for oxygen, CO2, humidity, pressure, and temperature. A detailed software program was also written to help efficiently carry out the parameters set out for use.

**Results:** The equipment is capable of automatically regulating the inner environment based on the set parameters. The combined cluster of regulators, efficiently controlled the oxygen levels, CO2 levels, and humidity levels within the experimental parameters.

**Conclusion:** This equipment thus acts as one of the easily reproducible simple assembly units with the integration of complex parameters, thus monitoring and controlling the inner environment with high precision.

## 1) Introduction

Hypoxia, a condition in which the body is deprived of adequate oxygen delivery, has been extensively researched due to its significant impact on human health and performance (Luo et al 2022). From the physiological impacts of high-altitude exposure to the therapeutic implications of low-oxygen conditions, hypoxia has proven to be an important area of research in a variety of fields (Chen et al 2020, Navarette-opazo et al 2014). In recent years, there has been increased interest in the use of hypoxic environment-stimulating devices, which aim to duplicate high-altitude exposure circumstances in a controlled and safe manner(Mallet et al 2023, Burtscher et al 2022). These devices have shown promise in improving cardiovascular health (Abe et al 2017), enhancing sports performance, and treating medical problems such as sleep apnea (Regev et al 2022), neurological dysfunction (Li et al 2023) and chronic obstructive lung disease (Timon et al 2022), and also extensively employed in stemcell research(Ito et al 2015, Lee et al 2020). Commercial hypoxia/hyperoxia chambers with environmental monitoring are available, but they are highly expensive and need proprietary software that costs a lot of money to subscribe to or buy. Because they are frequently created for single animal systems and are unable to withstand prolonged or irregular hypoxic exposure, these systems have limitations.

Conventional hypoxia chambers, on the other hand, resemble laminar airflow chambers in design and are huge chambers with an inlet flow of the desired hypoxic concentration of gasses(Luo et al 2022). Accounting for frequent gas exchanges, these instruments often fail to monitor carbon dioxide levels, and humidity levels, which directly influence animal comfort, and thus the experimental outcomes. These chambers’ huge size has drawbacks, including an inconsistent hypoxic environment and significant operating and maintenance costs (Saxena et al 2020).

In spite of this, the current technology for the simulation of hypoxia does have some drawbacks. Each experimental run necessitates a large amount of gas, the gas inside the chamber is not homogeneous, anaerobic gas packs cannot adjust gas concentrations, and real-time imaging is not possible

Our team has designed a low-cost hypoxia chamber for experimental rats that addresses the limitations of previous designs. The chamber is designed to accommodate the maximum number of animals while monitoring various parameters, including oxygen and CO2 levels, temperature, humidity, and air pressure, to maintain homeostasis. We incorporated several parts, such as a CO2 adsorber, dehumidifier, automated N2 gas, and O2 gas inlets, to ensure the setup provides optimized environmental conditions. All of the hardware was procured locally, and we did not use any custom-made electronics in the procedure, making it easy to replicate in other labs around the world. We developed the sketch based on C++ using Arduino and made it open-source, allowing for easy access to the design. By combining all of these elements, we were able to achieve a low-cost hypoxia chamber that met our requirements (Table-1). We therefore designed this study module, at a cheap cost and in an easily repeatable method, to solve the deficiencies from past research by monitoring and adjusting several factors, to preserve homeostasis.

**Table 1:**
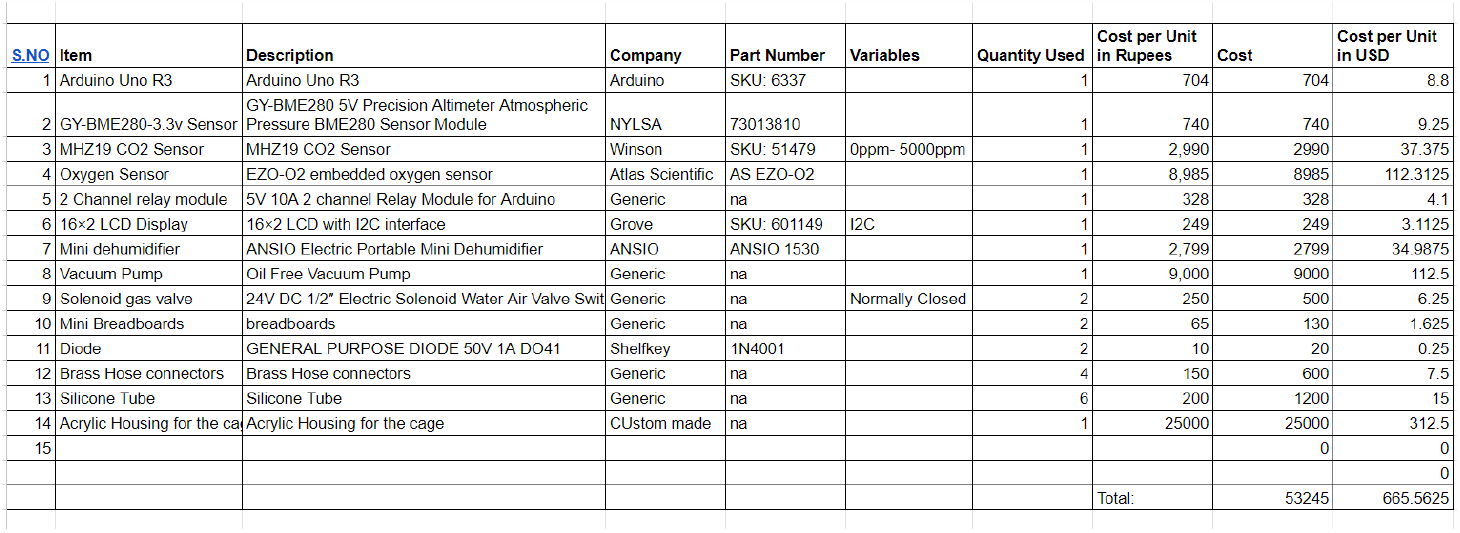
List of Materials used and their cost:

## 2) Methods

### 2.1) Hypoxia Chamber Fabrication

The hypoxic chamber, which measures 75 cm x 60 cm x 60 cm and is created in the shape of a cuboid, is made of 0.8 cm thick acrylic sheets that are properly glued together. The chamber is separated into three areas, with the animal cages being housed in the larger animal housing area, the electronic components being housed in the upper electronics housing area, and the CO2 adsorber and dehumidifier being housed in the lower area. The hypoxic chamber also contains a number of openings, including one on the bottom left side to collect inside air for the CO2 adsorber and four on the upper right side to house the N2 gas intake, O2 gas inlet, CO2 adsorber inlet, and power supply for electronics, respectively. The front of the chamber has two large doors that make loading and unloading the animal cages simple (Figure-1).

**Figure 1:**
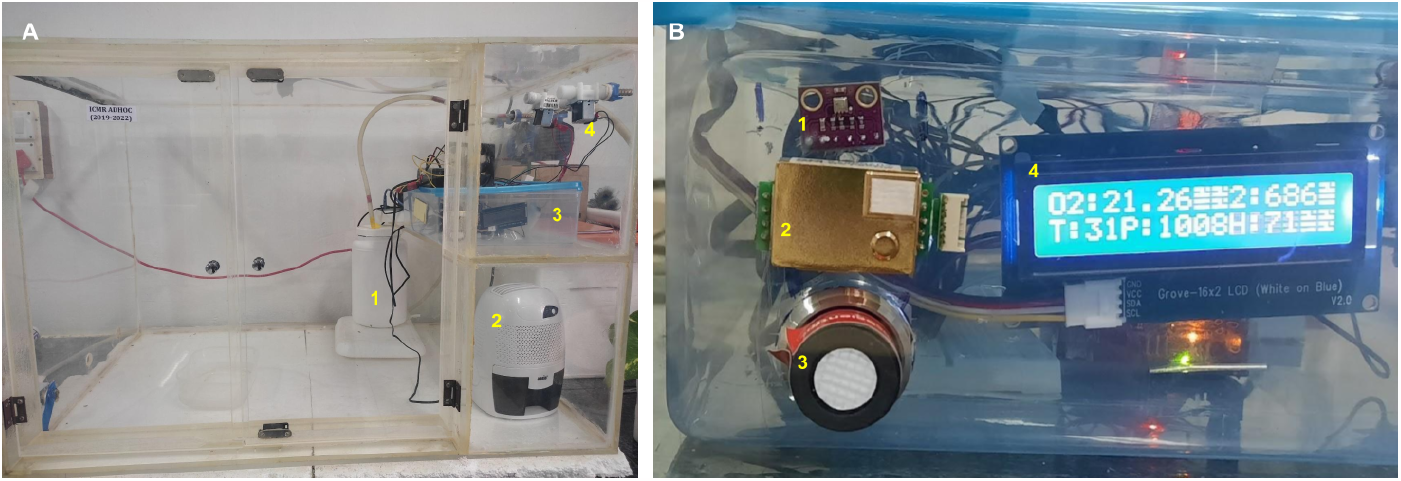
Hypoxia Chamber Module: (A) Acrylic-based physical chamber comprises A1- CO2 adsorber, A2- Dehumidifier, A3- Electronics module, and A4- Gas inlets. Close up view of A3 is shown in (B), comprising B1- BME280 sensor, B2- MH-Z19B CO2 sensor, B3- EZO-O2 sensor, and B4- 16x2 LCD display, showing O2, and CO2 levels in the upper line, while showing temperature, pressure, and humidity in the lower line.

### 2.2) Microcontroller Board

Due to its versatility and ease of use, Arduino Uno is one of the most extensively used boards in the Arduino family and hence utilized in this project. Six analog input pins on the Arduino Uno can read analog signals between 0 and 5 volts. For connecting with sensors that produce analog signals, these pins were used. The USB interface on the Arduino Uno is used for both power supply and programming. As a result, the board may be readily programmed using the user-friendly software known as the Arduino Integrated Development Environment (IDE).

### 2.3) Input devices (Sensors)

#### 2.3.1) Oxygen Gas Sensor

The Atlas Scientific EZO-O2 sensor was used to measure the partial pressure of oxygen through reduction. The sensor, a small fuel cell, can detect O2 levels between 0 and 420 ppt with a resolution of 0.01. The sensor can efficiently communicate with Arduino uno using I2C data protocols. The pin connections for this protocol are depicted in Figure 2.

**Figure 2:**
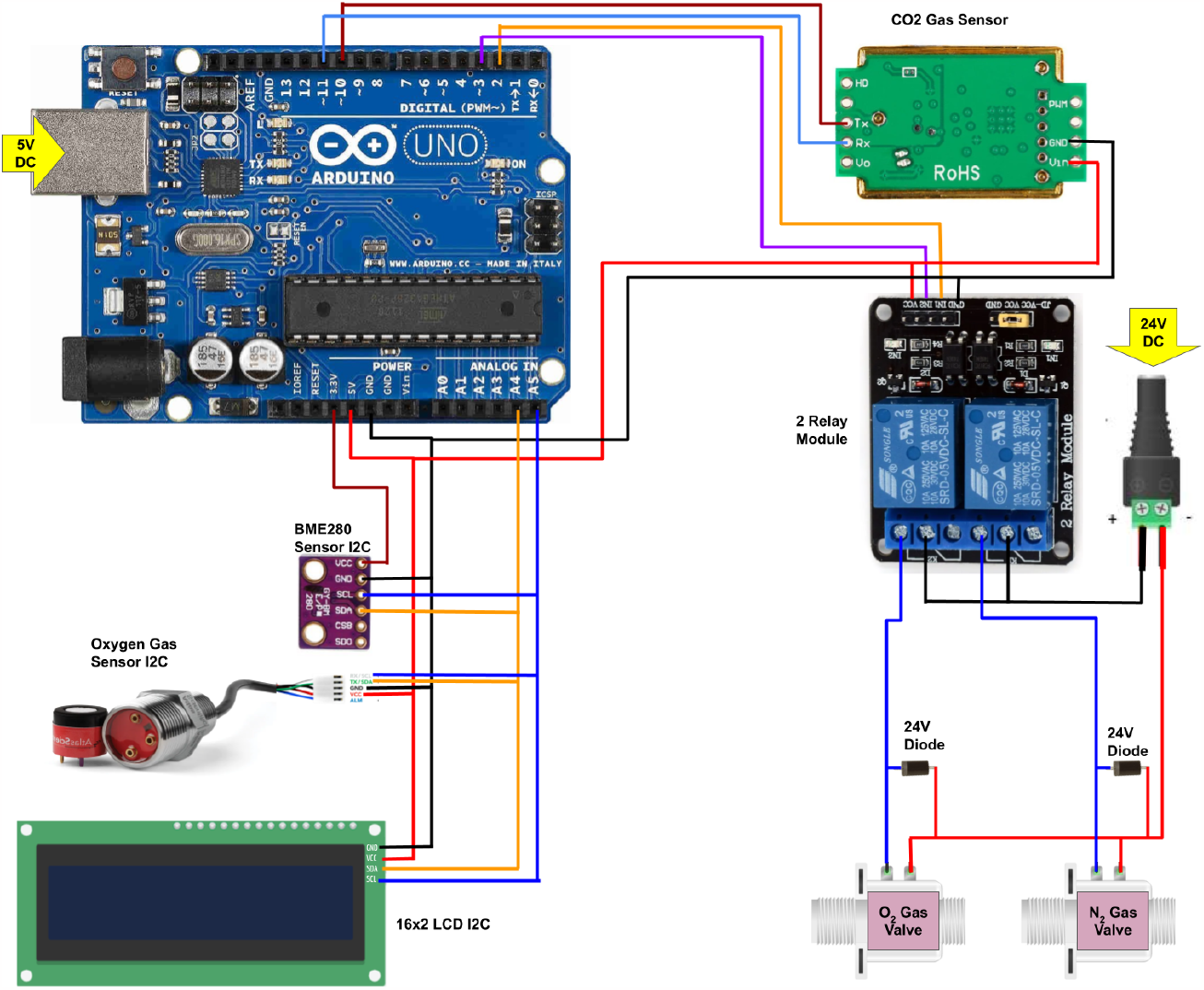
Wiring Diagram: The above diagram illustrates how the Arduino, sensors, LCD display, 2 relay module, and gas inlet valves are wired together.

#### 2.3.2) Carbon Dioxide Gas Sensor

The MH-Z19B NDIR infrared gas module is a small yet powerful CO2 sensor that is commonly used in research applications. It uses the non-dispersive infrared (NDIR) principle to detect the presence of carbon dioxide in the air with excellent selectivity and non-oxygen dependence. The sensor has a built-in temperature compensation feature that ensures accurate and reliable readings even in varying temperature conditions. With a detection range of 400∼5000ppm and an accuracy of ± (50ppm+5%reading value), the MH-Z19B NDIR infrared gas module is a reliable and accurate sensor that can be used in a range of research applications requiring CO2 measurement. The pin connections for this sensor are depicted in the figure-2, which also uses the UART protocol.

#### 2.3.3) BME 280 Sensor

The BME280 sensor is a powerful tool for researchers, as it combines digital humidity, pressure, and temperature measurements into a single device(Sari et al 2022). The BME280 is particularly effective at measuring humidity, with an impressive accuracy of ±3%, as well as barometric pressure with an absolute accuracy of ±1 hPa, and temperature with an accuracy of ±1.0°C. The pin connections for this protocol are depicted in the figure-2, which also uses the I2C protocol.

### 2.4) Output Devices

An electronic setup based on Arduino that had two main outputs were used to keep the experimental chamber in a state of homeostasis. A 16×2 LCD display with a low cost was the first output and showed the measured parameters’ current values. As a result, scientists could keep an eye on the circumstances inside the chamber and make any necessary adjustments. The N2 and O2 gas inlet valves were controlled separately by a 2-relay module, which was attached to the second output. To the appropriate gas cylinders outside the chamber, these gas inlets were connected. The relay module managed when and how much nitrogen and oxygen gas entered the chamber using a signal from the Arduino board. This made it possible for the researchers to keep the ideal gas mixture for their experiment and guarantee that homeostasis persisted throughout the experiment. Overall, this setup gave us a dependable, affordable way to keep the chamber’s experimental conditions where we wanted them.

### 2.5) CO2 Adsorber & Dehumidifier

In order to control the severe increase in CO2 release and humidity levels inside the hypoxic chamber designed for small animals like rats, two separate apparatuses were used. A custom CO2 scrubber was used to control the CO2 levels within the chamber. The scrubber consisted of a large plastic jar filled with medical-grade color-changing soda lime. Air from inside the chamber was pumped into the jar through an inlet and then circulated back into the chamber through an outlet. This allowed the soda lime to absorb the CO2 and prevent its accumulation within the chamber. Also, the inlet for the adsorber is placed on the chamber wall opposite to the chamber wall where the CO2 adsorber outlet is present, thus enabling cross-chamber air flow to achieve a homogeneous interior environment. Additionally, an electronic dehumidifier was used to bring down the increasing humidity levels during the hypoxic treatment procedures. Together, these apparatuses provided an effective solution for maintaining the desired environmental conditions within the hypoxic chamber and ensured that the experiment was conducted under controlled conditions that were conducive to the health and well-being of the animals.

### 2.7) Sketch

In this study, the program for the Arduino microcontroller was a crucial component of the experimental setup. The program was designed to collect input from various sensors, including the EZO-O2 sensor, the BME280, and the MH-Z19B CO2 sensor which were used to monitor the O2, temperature, humidity, and CO2 levels inside the hypoxic chamber. Based on the input from these sensors, the program executed clause-based outputs to control the gas inlets connected to appropriate gas cylinders outside the chamber. This allowed for precise control over the amount and timing of nitrogen and oxygen gas entering the chamber, which was critical for maintaining the homeostatic conditions required for the experiment. Additionally, the software was programmed to output the sensor readings onto the 16×2 LCD, allowing for easy real-time observation of the environmental conditions inside the chamber. Overall, the program for the Arduino microcontroller ensured that the experimental conditions were monitored and controlled accurately and effectively throughout the course of the study. The Arduino sketch is attached as a supplementary to the article, and also available in github (https://github.com/RoshanAasish/Hypoxia-Chamber).

### 2.8) Equipment Validation

In this study, Wistar albino rats weighing between 250-300g were used for the trial studies. The rats were housed in various numbers ranging from 1 to 9 at a time to observe the capabilities of the hypoxic chamber. The primary experiment involved exposing the animals to intermittent hypoxia at 11-14% oxygen levels for 4 hours per day. During this period, various parameters such as the oxygen and carbon dioxide levels, temperature, and humidity inside the chamber were monitored continuously using the electronic setup described earlier. All experiments were conducted in accordance with the approval of the Institutional Animal Ethics Committee (IAEC) (Ref No: BDU/IAEC/P07/2019, November 11, 2019), under the regulation of the Committee for the Purpose of Control and Supervision of Experiments on Animals (CPCSEA), India. The results of these analyses are discussed in detail in the subsequent sections of this paper.

## 3) Results

### 3.1) Hypoxia Chamber Validation

The initial experimental conditions were achieved by purging 100% N2 gas into the chamber, which took around 4 minutes to bring the O2 level from atmospheric 21% to experimental 11%. Therefore, a short single-step purging was sufficient to bring the atmospheric conditions to experimental conditions. The average temperature conditions during the initiation of the experiment were around 32 degrees Celsius, and the average air pressure during initial conditions was around 1005hPa to 1008hPa. The average carbon dioxide levels ranged around 650 ppm, and the humidity was around 50% during the initiation of the experiment.

### 3.2) Hypoxia chamber efficiently maintains oxygen level

After the first purge, the oxygen level in the chamber decreased to around 11% from 21%. The system was programmed to cut off the nitrogen pump once the O2 level reached 12%. Additionally, when the O2 level reached above 14%, the system opened the N2 gas inlet to bring back the oxygen level to 11%. This ensured that the O2 levels inside the chamber were always maintained within the 11-14% range, which was sufficient to induce proper intermittent hypoxic conditions. The fluctuations in oxygen levels were shown in the figure-3. However, a single purge was not enough to maintain the oxygen level within the range for the entire 4-hour experiment. On average, 2-3 N2 gas purges were needed to maintain the oxygen at experimental levels. Since the system was automated, no personal opening and closing of valves was required.

**Figure 3:**
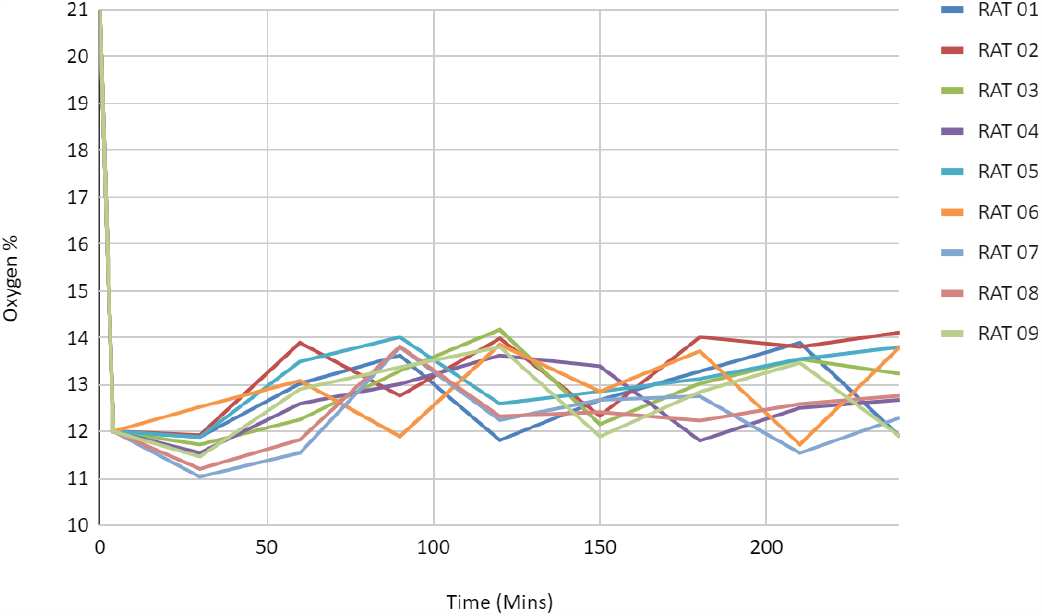
Hypoxia chamber efficiently maintains oxygen level: When the O2 level reaches 12%, the system is set to shut off the nitrogen pump and open the N2 gas input, guaranteeing that the O2 levels are kept between 11 and 14%.

### 3.3) CO2 adsorber is crucial in regulating CO2 levels within homeostatic conditions

At the beginning of the experiment, the atmospheric CO2 levels were approximately 650 ppm, but after initiation, these levels increased steadily to toxic levels exceeding 5000 ppm. The rate of CO2 accumulation is proportional to the number of animals exposed to the experiment simultaneously. To maintain CO2 levels below 5000 ppm, the CO2 adsorber played a critical role. The population-dependent increase in CO2 levels and their regulation by the CO2 adsorber is depicted in the figure-4. The equipment was efficient enough to maintain CO2 levels below 1500 ppm during a single animal experiment, and even below 4000 ppm during experiments involving up to 9 animals. Therefore, the equipment was capable of regulating CO2 levels as needed, even in a highly populous environment with 9 adult rats.

**Figure 4:**
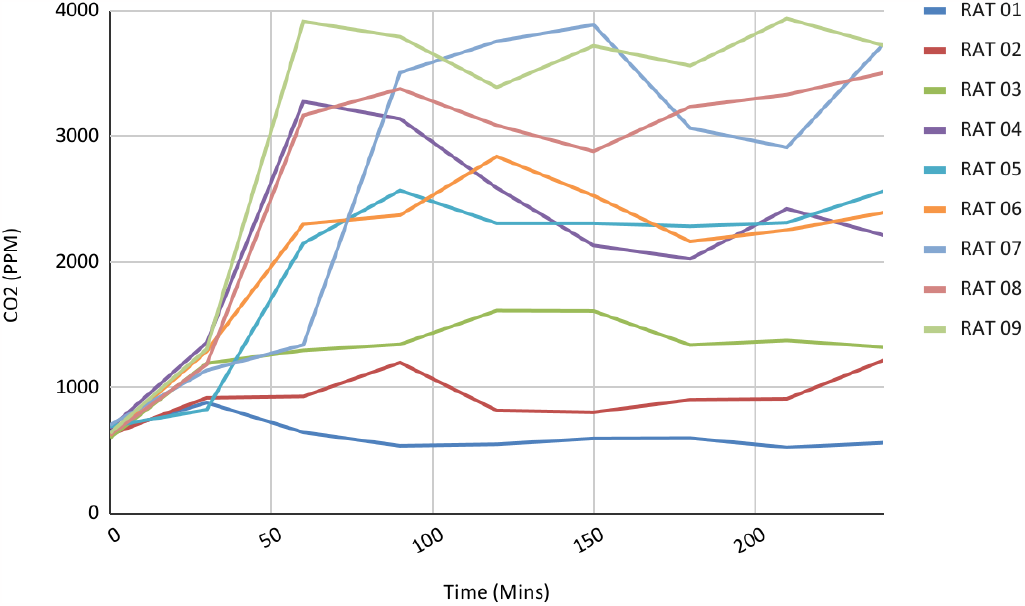
CO2 adsorber is crucial in regulating CO2 levels within homeostatic conditions: The atmospheric CO2 concentration was 650 ppm at the start of the experiment, but gradually increased to around 4000 ppm, with the rate of accumulation proportional to the number of animals exposed. Co2 adsorber efficiently controlled CO2 levels under 4000ppm, despite the number of animals inside reaching 9.

### 3.4) Dehumidifiers effectively lower the excessive humidity brought on by hypoxic exposure

At the start of the experiment, the humidity level was at 45%. However, without a dehumidifier, the humidity levels steadily increased, reaching 60% mid-way through the experiment and even rising to 70% later on. The Humidifier efficiently controlled humidity levels below 50% throughout the experiment (Figure-5).

**Figure 5:**
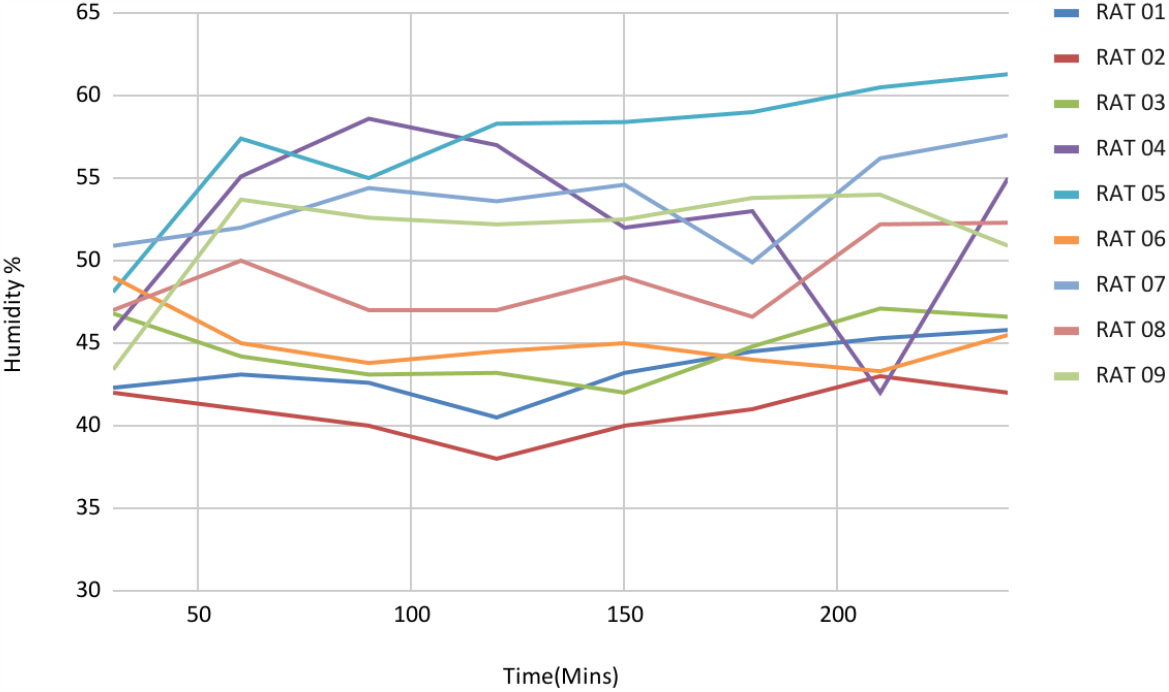
Dehumidifier efficiently lowers the excessive humidity brought in by hypoxic exposure: Throughout the experiment, the humidifier successfully maintained humidity levels around 50%.

## 4) Discussion

This paper presents the design and validation of a low-cost portable oxygen sensor platform for monitoring and logging oxygen tension during hypoxia-based studies. The platform is easily assembled by researchers and can be adapted to accommodate different in-vivo or in-vitro models. To prioritize easy visualization of animal cages and facilitate cleaning, acrylic sheets were used. However, this decision increased the project cost, as alternatives could have been chosen to reduce the expense of the acrylic sheets, which amounted to approximately $300. Compartmentalizing the chambers allowed for easier wiring and connection of gas flow equipment, resulting in a tidier and more navigable chamber interior.

The electronic setup in the hypoxic chamber utilized an Arduino Uno microcontroller board. This choice was based on the availability of extensive do-it-yourself (DIY) projects online, such as those found on GitHub and Instructables. These existing projects greatly assisted in circuit design and the development of the Arduino sketch. While there is a wide range of gas sensors available worldwide, each with its own advantages and associated costs, our options were limited to procuring from online sites available in India. Many international companies were uninterested in shipping products with a low net value, as the shipping costs exceeded the product costs. From the available options, we selected the Atlas Scientific EZO-O2 sensor for measuring oxygen levels, the MH-Z19B NDIR infrared gas module for measuring carbon dioxide levels, and the BME280 sensor for measuring humidity, pressure, and temperature. These sensors were chosen for their accuracy, reliability, and compatibility with UART or I2C data protocols for communication with the Arduino Uno microcontroller board.

To maintain the oxygen level within the desired range of 11-14% throughout the 4-hour experiment, the hypoxic chamber was equipped with automatic monitoring and management of N2 and O2 gas inlets. Fluctuations in oxygen levels (Figure 3) were achieved through N2 gas purges. By minimizing the frequency of gas purges, we significantly reduced nitrogen gas usage, allowing the operation of the hypoxic chamber using N2 gas cylinders. This approach eliminated the need for an expensive gas mixer, which is often overlooked by others.

Rodents exhibit high metabolic activity and exhale concentrated amounts of CO2 in a short period of time. Efficient CO2 regulation was crucial when accommodating more than 6 rats in the small confined space of the hypoxic chamber. Taking inspiration from CO2 adsorber systems used in human surgery theaters during anesthesia, we developed our own CO2 adsorber to effectively regulate CO2 levels. The Occupational Safety and Health Administration (OSHA) Permissible Exposure Limit (PEL) and the American Conference of Governmental Industrial Hygienists (ACGIH) Threshold Limit Value (TLV) for 8-hour exposure state that CO2 levels should not exceed 5000ppm (Fan et al 2023, Azuma et al 2018). Our equipment successfully maintained CO2 levels below 1500 ppm during single-animal experiments and below 4000 ppm during experiments involving up to 9 animals. Figure 3 in our data shows that CO2 accumulation is proportional to the number of animals. Additionally, a dehumidifier was employed to maintain humidity levels below 50% throughout the experiment, as high humidity levels have been reported to induce oxidative stress-related inflammation in rats (Mertoglu et al 2020). Finally, the custom CO2 scrubber and dehumidifier played a crucial role in maintaining the experimental conditions within the hypoxic chamber

The output devices used in the electronic setup included a 16×2 LCD display for presenting measured parameter values and a 2-relay module for controlling the nitrogen and oxygen gas inlet valves. These output devices enabled researchers to monitor experimental conditions inside the chamber and make necessary adjustments to ensure the desired hypoxic conditions for the animals.

### Future Perspectives

Though this model was efficient in monitoring multiple parameters, it lacked the ability to transmit the data through wifi to the experimenter’s computer or handheld devices, This would efficiently enable the experimenter to multitask while the experiment is ongoing. Also, an efficient Graphic user interface will easily enable altering the experimental conditions easier, rather than changing them by modifying the Arduino sketch. Also, a sophisticated model of CO2 adsorber and dehumidifier which feedback-based regulation will greatly improve the efficiency of the current system.

Although the current model demonstrated proficiency in monitoring various parameters, it exhibited limitations in transmitting data via wifi to the experimenter’s computer or handheld devices. This functionality would greatly facilitate multitasking for the experimenter during ongoing experiments. Additionally, the implementation of an efficient graphic user interface would simplify the process of modifying experimental conditions, eliminating the need for manually modifying the Arduino sketch. This enhancement would streamline the experimental workflow. Furthermore, the integration of a sophisticated CO2 adsorber and dehumidifier, incorporating feedback-based regulation, holds immense potential for enhancing the efficiency of the existing system. This advancement would significantly contribute to the overall performance and optimization of the system.

## Conclusion

Overall, the hypoxic chamber described in this research paper provides researchers with a reliable and accurate tool for studying the effects of low oxygen levels on small animals. The use of high-quality electronic components and the custom CO2 scrubber and dehumidifier help to ensure that the experimental conditions are maintained throughout the experiment, providing researchers with reliable and accurate data for their studies. This low-cost platform is particularly useful for researchers who cannot afford expensive instruments for hypoxia-based studies.

## Contributions

Conceptualization: SAR, AM; design construction and data acquisition: SAR; analysis and interpretation of data: SAR, SKJ, MK, AM; writing–original draft preparation: SAR; writing–review and editing: SAR, SKJ, MK, AM; critical revision of the manuscript for intellectual content: SKJ, MK, AM; supervision: AM

## Acknowledgments

The authors would like to thank Aishwarya Suresh for proofreading the manuscript and for helpful suggestions.

## Data Availability

The data supporting the findings of this study are available within the article and its supplementary material.

