## Supplementary file-1 for "Design and Evaluation of an Affordable Hypoxic Chamber with Comprehensive Environmental Control for Research Applications"

**Article title**

**Arduino Sketch:**

#include <Adafruit_BME280.h>

#include <Ezo_i2c.h> //include the EZO I2C library from https://github.com/Atlas-Scientific/Ezo_I2c_lib

#include <Wire.h>	//include arduinos i2c library

#include <rgb_lcd.h> //include lcd library

#include "MHZ19.h"

#include <SoftwareSerial.h> // Remove if using HardwareSerial or Arduino package without SoftwareSerial support

#define RX_PIN 10	// Rx pin which the MHZ19 Tx pin is attached to

#define TX_PIN 11	// Tx pin which the MHZ19 Rx pin is attached to

#define BAUDRATE 9600 // Device to MH-Z19 Serial baudrate (should not be changed)

MHZ19 myMHZ19; // Constructor for library

SoftwareSerial mySerial(RX_PIN, TX_PIN); // (Uno example) create device to MH-Z19 serial

//HardwareSerial mySerial(1); // (ESP32 Example) create device to MH-Z19 serial

unsigned long getDataTimer = 0;

Ezo_board OXY = Ezo_board(108, "OXY"); //create a OXY circuit object, who's address is 108 and name is "OXY"

rgb_lcd lcd;

bool reading_request_phase = true; //selects our phase

uint32_t next_poll_time = 0; //holds the next time we receive a response, in milliseconds

const unsigned int response_delay = 1000; //how long we wait to receive a response, in milliseconds

String Oxystring = ""; //a string to hold the data from the Atlas Scientific product

float O2; //used to hold a float number that is the o2

float CO2ppm; // used to hold float number that is the CO2

unsigned long ms_from_start =0;

unsigned long ms_previous_read_N2Valve =0;

unsigned long N2Valve_interval =10000;

unsigned long ms_previous_read_O2Valve =0;

unsigned long O2Valve_interval =5000;

unsigned long ms_previous_read_Vacuum =0;

unsigned long Vacuum_interval =3500;

const int solenoidPinN2 = 2; // Solenoid valve for N2 gas

const int solenoidPinO2 = 3; // Solenoid Valve for O2 Gas

const int vacuumPin = 13; // Vacuum pump

float temperature;

float humidity;

float pressure;

float altitude;

float const ALTITUDE = 62.0; // Altitude at my location in meters

float const SEA_LEVEL_PRESSURE = 1013.25; // Pressure at sea level

Adafruit_BME280 bme; // I2C

void setup() {

Wire.begin(); //start the I2C

lcd.begin(16,2);

Serial.begin(9600); //start the serial communication to the computer

mySerial.begin(BAUDRATE); // (Uno example) device to MH-Z19 serial start

//mySerial.begin(BAUDRATE, SERIAL_8N1, RX_PIN, TX_PIN); // (ESP32 Example) device to MH-Z19 serial start

myMHZ19.begin(mySerial); // *Serial(Stream) refence must be passed to library begin().

myMHZ19.autoCalibration(); // Turn auto calibration ON (OFF autoCalibration(false))

pinMode(solenoidPinN2, OUTPUT); //N2 gas

pinMode(solenoidPinO2, OUTPUT); //O2 gas

pinMode(vacuumPin, OUTPUT); //O2 gas

bool status;

// default settings

status = bme.begin(0x76); // The I2C address of the sensor is 0x76

}

void loop() {

getPressure(); // Get sensor data and print to LCD

getHumidity();

{

if (reading_request_phase) {	//if were in the phase where we ask for a reading

//send a read command. we use this command instead of PH.send_cmd("R");

//to let the library know to parse the reading

OXY.send_read_cmd();

next_poll_time = millis() + response_delay; //set when the response will arrive

reading_request_phase = false; //switch to the receiving phase

}

else {	//if were in the receiving phase

if (millis() >= next_poll_time) { //and its time to get the response

receive_reading(OXY); //get the reading from the OXY circuit

Serial.print(" ");

reading_request_phase = true; //switch back to asking for readings

}

}

}

{	if (millis() - getDataTimer >= 2000)

{

String CO2;

/* note: getCO2() default is command "CO2 Unlimited". This returns the correct CO2 reading even

if below background CO2 levels or above range (useful to validate sensor). You can use the

usual documented command with getCO2(false) */

CO2 = myMHZ19.getCO2(); // Request CO2 (as ppm)

CO2ppm = CO2.toFloat(); //convert the string to a float so it can be evaluated by the Arduino

Serial.print("CO2 (ppm): ");

Serial.println(CO2);

lcd.setCursor(8,0);

lcd.print("CO2:");

lcd.println(CO2);

int8_t Temp;

Temp = myMHZ19.getTemperature(); // Request Temperature (as Celsius)

Serial.print("Temperature (C): ");

Serial.println(Temp);

lcd.setCursor(0,1);

lcd.print("T:");

lcd.println(Temp);

getDataTimer = millis();

}

}

{

ms_from_start =millis();

if (ms_from_start-ms_previous_read_N2Valve> N2Valve_interval) {

ms_previous_read_N2Valve = ms_from_start;

if (O2 < 12.50) {

digitalWrite(solenoidPinN2, HIGH); //switch solenoid OFF

Serial.print("N2Valve is OFF");

} else if (O2 > 14.00) {

digitalWrite(solenoidPinN2, LOW); //switch solenoid ON

Serial.print("N2Valve is ON");

;

}}

if (ms_from_start-ms_previous_read_O2Valve> O2Valve_interval) {

ms_previous_read_O2Valve = ms_from_start;

if (O2 > 14.00) {

digitalWrite(solenoidPinO2, HIGH); //switch solenoid OFF

Serial.print("O2Valve is OFF");

} else if (O2 < 11.00) {

digitalWrite(solenoidPinO2, LOW); //switch solenoid ON

Serial.print("O2Valve is ON");}}

if (ms_from_start-ms_previous_read_Vacuum> Vacuum_interval) {

ms_previous_read_Vacuum = ms_from_start;

if (CO2ppm< 2000) {

digitalWrite(vacuumPin, HIGH); //switch vacuum OFF

Serial.print("Vacuum is OFF");

} else if (CO2ppm> 2000) {

digitalWrite(vacuumPin, LOW); //switch vacuum ON

Serial.print("Vacuum is ON");}

}

}

}

void receive_reading(Ezo_board &Sensor) { // function to decode the reading after the read command was issued

Serial.print(Sensor.get_name()); Serial.print(": "); // print the name of the circuit getting the reading

Sensor.receive_read_cmd(); //get the response data and put it into the [Sensor].reading variable if successful

switch (Sensor.get_error()) {	//switch case based on what the response code is.

case Ezo_board::SUCCESS:

Serial.print(Sensor.get_last_received_reading()); //the command was successful, print the reading

Oxystring = Sensor.get_last_received_reading();

O2 = Oxystring.toFloat(); //convert the string to a float so it can be evaluated by the Arduino

lcd.setCursor(0,0);

lcd.print("O2:");

lcd.println(O2);

break;

case Ezo_board::FAIL:

Serial.print("Failed "); //means the command has failed.

break;

case Ezo_board::NOT_READY:

Serial.print("Pending "); //the command has not yet been finished calculating.

break;

case Ezo_board::NO_DATA:

Serial.print("No Data "); //the sensor has no data to send.

break;

}

}

//===============================================================================

// getHumidity - Subroutine to get and print humidity

//===============================================================================

void getHumidity()

{

humidity = bme.readHumidity();

// String humidityString = String(humidity, 0);

lcd.setCursor(10, 1);

lcd.print("H:");

lcd.println(humidity);

}

//===============================================================================

// getPressure - Subroutine to get and print pressure

//===============================================================================

void getPressure()

{

pressure = bme.readPressure();

pressure = bme.seaLevelForAltitude(ALTITUDE, pressure);

//pressure = pressure / 3386.39; // Convert hPa to in/Hg

lcd.setCursor(4, 1);

lcd.print("P:");

String pressureString = String(pressure, 0);

lcd.println(pressureString);

}
